# Optimizing crop clustering to minimize pathogen invasion in agriculture

**DOI:** 10.1101/2025.03.07.642019

**Authors:** Yevhen F. Suprunenko, Christopher A. Gilligan

## Abstract

**Context:** The initial rate of pathogen invasion in crops is influenced by the spatial clustering of susceptible crops and the characteristics of pathogen dispersal. Previous studies have shown that various degrees of crop clustering can effectively reduce this invasion rate. However, the optimal degrees of clustering that minimize pathogen invasion have not previously been identified.

**Objectives:** This study aims to determine analytically the range of crop clustering that minimizes the initial rate of pathogen invasion.

**Methods:** We studied artificial agricultural landscapes with crop areas arranged in identical square clusters on a regular square lattice. For pathogen dispersal, we used several common dispersal kernels, including Gaussian, negative exponential, and power-law. The optimal degree of clustering, defined by cluster size and separation distance, was calculated using a new analytical approximation for the pathogen invasion rate, which showed strong agreement with computer simulations. Additionally, we analysed a realistic cassava landscape at risk of invasion by cassava brown streak virus.

**Results:** We identified a range of optimal cluster sizes and corresponding separation distances that minimize pathogen invasion rates for various dispersal kernels and landscapes with clusters of crop fields arranged on a regular square lattice. The methods can be extended to other geometrical configurations, such as long narrow fields. Using a cassava landscape as an example, we show how optimal crop clustering strategies can be derived to mitigate the potential invasion of cassava brown streak virus.

**Conclusion:** The methods provides analytical insights that can help farmers and agricultural planners to optimize the spatial structure of agricultural landscapes to minimize initial pathogen invasion rates.

## 1. Introduction

One of the key factors that determine how quickly a crop pathogen (or pest) invades and spreads across an agricultural landscape is the relationship between the pathogen’s dispersal and the spatial distribution of susceptible crops [1–13]. The rate of pathogen invasion can be slowed by adjusting the spatial configuration of host distribution relative to pathogen dispersal [1,5,8,9,11,12,14,15]. This can be achieved by reducing host aggregation [16–21], growing resistant varieties [22], or applying fungicides [7,23].

Two simple measures of pathogen invasion are the infection rate, *r*, which denotes the early exponential growth rate of the number of infected fields at the onset of an epidemic, and the basic reproduction number *R*_0_. The latter is defined as *R*_0_ = *r*/*μ*, where 1/*μ* represents the average infectious period of infected hosts.

Insights from theoretical [1,5,11,12] and experimental [8,9] studies indicate that pathogen invasion slows, and both *r* and *R*_0_ decline, as the host landscape becomes more dispersed (holding all other factors constant). While this might suggest that the minimal values of *r* and *R*_0_ would occur when fields of susceptible crops are evenly dispersed, recent theoretical work [24] has shown that *r* and *R*_0_ can be minimized in agricultural landscapes with non-uniform spatial configurations. In these configurations, fields with susceptible crops can be aggregated into clusters (i.e. clustered) up to a threshold size.

Suprunenko et al. (2025) [24] illustrated this process using artificial landscapes in which a fixed area of susceptible crops was distributed in clusters on a regular square lattice (figure 1a). As the total crop area remained constant, the number of clusters (*N*_*clusters*_) decreased as cluster size increased. The highest degree of clustering (i.e. the smallest *N*_*clusters*_) associated with the minimum *r* can be characterised by a threshold cluster size of width, 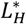, and separation distance between clusters, 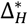 (cf figure 1b).

**Fig. 1.**
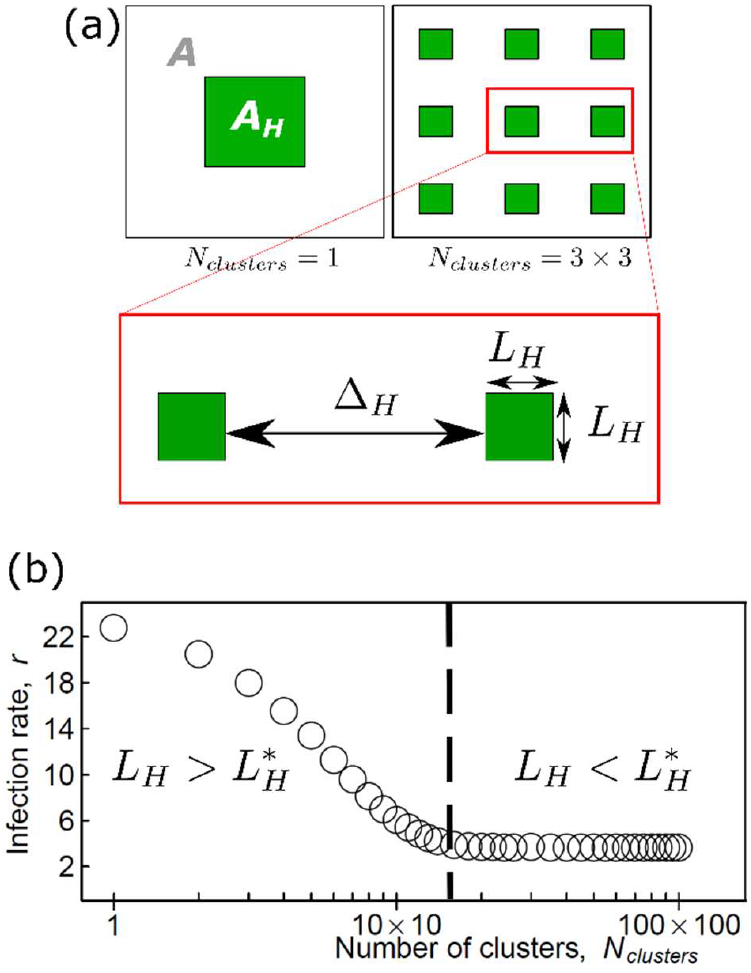
**(a)** An illustration of a crop area, *A*_*H*_ (green), aggregated into one cluster (*N*_*clusters*_ = 1) or nine clusters (*N*_*clusters*_ = 3 × 3), placed on a regular square grid within an agricultural landscape, *A*, where arbitrarily *A*_*H*_*A*^-1^ = 0.15. The degree of clustering (aggregation) is characterised by the number of clusters, *N*_*clusters*_, which determines the size (*L*_*H*_) of a square cluster and the separation distance (Δ_*H*_) between clusters. **(b)** Open circles indicate the infection rate *r* (the initial growth rate of the number of infected fields) estimated from simulations of an individual-based epidemic model initiated by a single randomly infected field within the landscapes of the type shown in Figure 1a ([24]; Supplementary Note 1). Reconfiguring the crop area *A*_*H*_ into clusters of size *L*_*H*_ smaller than the threshold size 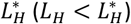 reduces the infection rate from its maximum value (corresponding to *N*_*clusters*_ = 1) to its minimum value (corresponding to 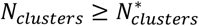, where 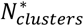 is the threshold number of clusters). Therefore, the infection rate of susceptible agricultural crop fields at the start of an epidemic can be minimized through various spatial reconfigurations of the landscape. The illustration is a modified version of figure 1 in Ref. [24]. Computer codes for all figures in the current paper are available from Figshare [25].

Understanding these threshold values, 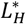 and 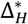, would be invaluable for agricultural planners in minimizing the risk of pathogen (or pest) invasion, given known dispersal characteristics.

Computer simulation of epidemic models on simulated landscapes can, in principle, be used to estimate the threshold size 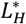 and separation distance 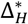 for a given pathogen dispersal kernel [15,20,21,26,27]. This involves analysing epidemics on host landscapes with many different spatial configurations to identify the degrees of host clustering that minimize *r* (or *R*_0_). However, constructing realistic epidemic spread models for each new pest or pathogen can be challenging, and considering numerous spatial configurations is computationally demanding and potentially inefficient.

Here, we explore an alternative approach to identify landscapes that minimise the initial rates of pathogen invasion using an analytical approximation for the initial infection rate, *r*. We test this analytical approximation against empirical infection rates derived from computer simulations of an individual-based model (IBM) for pathogen invasion and spread through a host landscape. Using the analytical approximation for *r*, we derive values for the threshold size 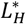 and separation distance 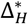 for a range of generic host landscapes and commonly-used dispersal kernels, including Gaussian, negative exponential, and power-law kernels, among others. We illustrate a potential application of this approach, motivated by the spread of cassava brown streak virus (CBSV), which threatens cassava production in sub-Saharan Africa [28–30]. Using a dispersal kernel for CBSV estimated by Godding *et al*. [31] and a section of a cassava landscape provided by Szyniszewska (2020) [32], we present examples of maps showing estimates of local threshold sizes and separation distances for clusters of cassava fields that minimise infection rate.

## 2. Methods

### 2.1 Mathematical approximation for infection rate

We consider a susceptible-infected (SI) epidemiological compartmental individual-based model (IBM) for the invasion and spread of a pathogen through an agricultural landscape partially occupied by susceptible host crops. An analytical expression for the infection rate, *r*, for this IBM has been variously derived in several studies, including those by Bolker (1999)[1], North and Godfray (2017)[12], van den Bosch et al. (2024)[33] and others [11,24,34,35]. For example, consider the approximation from Suprunenko et al. (2025)[24]:

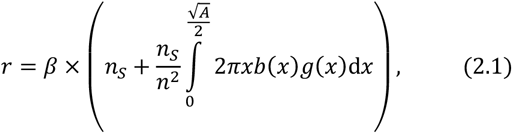

where *b*(*x*) is the pathogen dispersal kernel as a function of distance *x*, and *β* is the infection rate per contact density (i.e. the spatial density of susceptible hosts that an initial infected field can reach through the dispersal kernel *b*(*x*)). The host landscape is assumed to be a square plot of land with area *A* partially occupied by land containing a susceptible crop consisting of *N* sufficiently small, identical individual fields with area *A*_0_. Each field is treated as an individual ‘host’, described as a point-like object located at the centre of a field. The initial densities *n*_*S*_ = (*N* − 1)*A*^-1^ and *n* = *NA*^-1^ are the densities of susceptible hosts and all hosts respectively. The total area occupied by hosts is *A*_*H*_, i.e. *A*_*H*_ = *A*_0_*N*, where *A*_*H*_ ≤ *A*. The function, *g*(*x*), denotes the second order spatial cumulant (also known as a truncated correlation function or autocovariance, see e.g. [36,37], that characterises the spatial structure of a host landscape.

The expression inside the brackets in equation (2.1) represents the spatial density of contacts with susceptible hosts (i.e. fields) from a randomly located initial infected field. Similarly, van den Bosch et al. (2024)[33] described an equivalent product, *b*(*x*)*g*(*x*) (cf equation 2.1), as one that “describes how a pathogen ‘sees’ the host population around an infected host individual” [33]. Based on these insights, we construct an explicit expression for the spatial density of contacts to replace the expression in brackets in equation (2.1).

Our approach differs from conventional IBMs, which consider individual hosts as point-like entities and calculate the infection rate (2.1) based on their spatial distribution. In contrast, we explicitly consider individual hosts (i.e. fields) as square plots of cropped land with area *A*_0_, and estimate the infection rate using the spatial distribution of these land polygons. We denote the spatial density of fields (measured as the number of fields per area) at location ***x*** as *n*(***x***), defining 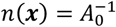 when a point ***x*** lies within a field, and *n*(***x***) = 0 when ***x*** is outside a field. A pathogen at location ***x*** can infect directly the density of fields *n*_b_(***x***), where

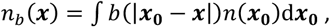

and the integration is over the entire landscape. Averaging *n*_*b*_(***x***) over all possible locations, ***x***, within the landscape leads to the quantity 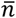,

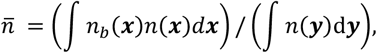

that can be interpreted as the spatially averaged density of contacts (with susceptible fields) that can result in infection from an initial randomly infected field. Therefore, the expression in brackets in equation (2.1) can be estimated by the density 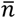, i.e.

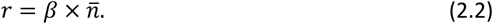

Further to this, we make the following two simplifying approximations.

#### Approximation 1

Here, we develop an approximation for equation (2.2) when there is a high degree of aggregation of the target crop within a landscape (i.e. when *N*_*clusters*_ is small):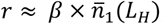, where 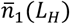 is derived as follows. We assume that the crop area is distributed into identical *L*_*H*_ × *L*_*H*_ square clusters on a regular square lattice, as illustrated in figure 1a. The total number of clusters is *N*_*clusters*_. We denote the spatial density of crops within a single cluster as *n*_1_(***x***), where 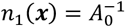 if 0 ≤ *x*_1_ ≤ *L*_H_ and 0 ≤ *x*_2_ ≤ *L*_*H*_ (where *x*_1_ and *x*_2_ are the coordinates in a two-dimensional Cartesian coordinate system), and *n*_1_(***x***) = 0 otherwise. Given a high degree of clustering, i.e. when all hosts are aggregated into a small number of large clusters, we estimate *r* on a single cluster and use *n*_1_(***x***) instead of *n*(***x***) in the expression (2.2), obtaining 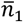 instead of 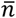, where

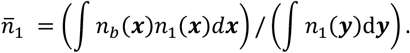

Using straightforward algebraic transformations, shown in the electronic supplementary material (Supplementary Note 2), the expression for 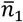 is simplified further:

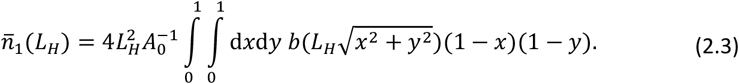

The quantity 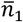 denotes the averaged density of fields within a single cluster that a randomly introduced infected field within that cluster can infect directly via a dispersal kernel *b*(*x*). The upper bound of the quantity 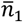 is 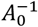, i.e. the density of crops within a single cluster.

#### Approximation 2

Here, we develop an approximation for equation (2.2) when there is a low degree of clustering in the host landscape: *r ≈ β* × *n*. We assume that the pathogen dispersal kernel, *b*(*x*), is sufficiently long-ranged relative to an individual field of area *A*_0_, i.e. the characteristic dispersal length, *a*, should be larger than the size of an individual field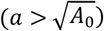. Consequently, the minimum value of the infection rate can be approximated as *r*_min_ = *β* × *n*, where *n* is the spatial density of all *N* fields of a host crop within the landscape with area *A, n* = *NA*^-1^. For a low degree of clustering, we therefore use *r ≈ r*_min_.

Finally, we combine expressions for the infection rate within a single cluster, 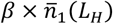 (approximation 1), and its minimal value, *β* × *n* (approximation 2) noting that the larger estimate should be used between approximations 1 and 2. Indeed, when the crop is highly aggregated within a landscape, the estimate of *r* is given by 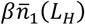 where 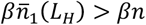; while for a low degree of crop aggregation the estimate of *r* is given by *βn* when 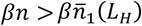. Therefore, using the maximum of two estimates, we use the following approximation for *r*, denoted as *r*_*approximate*_:

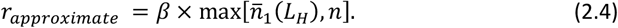

It is important to stress that this approximation is applicable to dispersal kernels, *b*(*x*), that are decreasing functions of distance *x*. For dispersal kernels that do not decrease monotonically with distance, such as inverse Gaussian (Wald) or lognormal dispersal kernels (see [38]) the infection rate does not necessarily decrease as the number of clusters (*N*_*clusters*_) increases.

### 2.2 Computer simulations of an individual-based model

To test the approximation derived above, we compare the performance of equation (2.4) with empirical rates for *r* at the start of an epidemic, originally derived from computer simulations of an individual-based model of pathogen invasion in an agricultural landscape [24]. For convenience, details of the individual-based model are summarised in Supplementary Note 1. Exploratory testing indicated that the transition of *r* from its maximum to its minimum value captured by *r*_*approximate*_ (equation 2.4) sufficiently well (see caption of figure 2); therefore, we proceed to use this approximation.

**Fig. 2.**
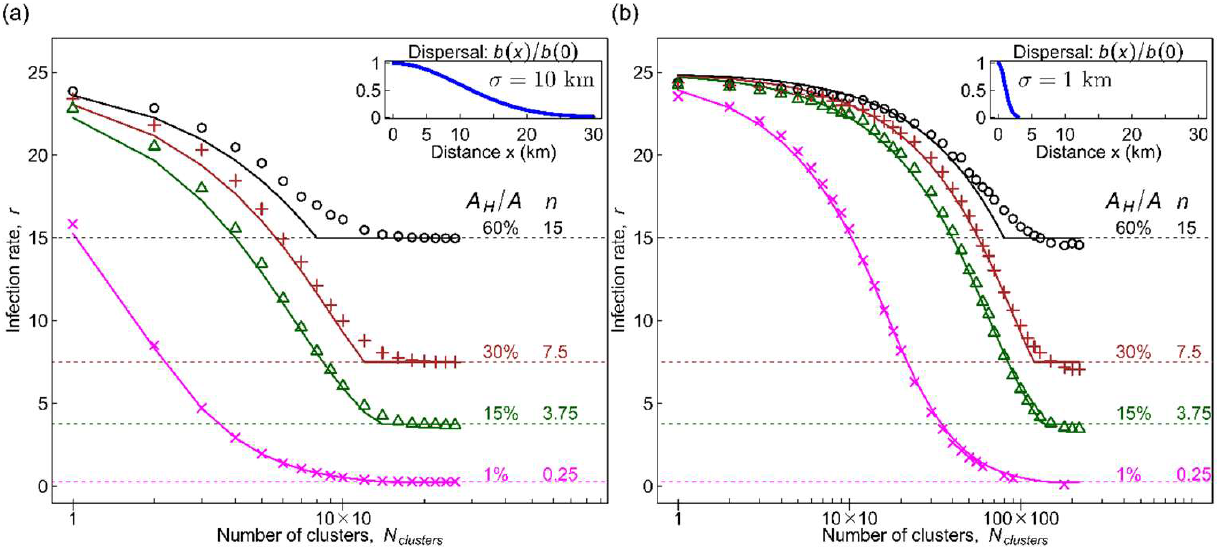
The behaviour of the analytically approximated infection rate, *r*_*approximate*_, (lines corresponding to equation 2.4) as a function of the degree of clustering of susceptible crops is compared with estimates of infection rate *r* (symbols) inferred from computer simulations of an individual-based model (see Methods). Results are shown for different landscapes and for two different dispersal kernels given by a Gaussian function with standard deviation: **(a)** *σ* = 10 km, and **(b)** *σ* = 1 km. The quantity *A*_*H*_*A*^-1^ denotes the fraction of the landscape occupied by susceptible fields; *n* = *NA*^-1^ is the spatial density of all *N* crop fields in the landscape with area *A*. Here, *β* = 1, and the area of a unit field is *A*_0_ = 0.04 km^2^; the upper bound of the rate *r* is given by 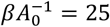. The transition of *r* from its maximum to minimum is captured by *r*_*approximate*_ reasonably well, as quantified by the following measures. In both panels, the maximum deviation max |*r* − *r*_*approximate*_| is 0.6, 0.9, 1.3 and 2.0 for *A*_*H*_/*A* = 1, 15, 30 and 60, respectively. These deviations correspond to 3.7%, 4.5%, 8.2% and 22% of the total range max(*r*) − min(*r*) for the same values of *A*_*H*_/*A*.

In figure 2 we used the data [39] from computer simulations of individual-based model of pathogen invasion and spread in agricultural landscapes that were taken from [24]. Here we test a different analytical approximation from [24] for identifying optimal clustering to reduce initial invasion rate, inspired by the original more computationally intensive method in [24] that uses spatial cumulants to assess the impact of general landscape structures.

### 2.3 Mathematical approximation for threshold size and separation distance

We define the threshold size 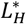 as the value of the argument of the function 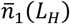 in equation (2.4) such that

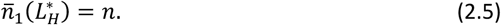

Using the definition *n* = *NA*^-1^ = *A*_*H*_(*AA*_0_)^-1^, the condition (2.5) can be re-written as

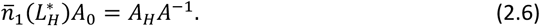

The right-hand side of equation (2.6) is a dimensionless parameter *A*_*H*_*A*^-1^ denoting the fraction of the landscape occupied by susceptible fields. For the most typical dispersal kernels,*b*(*x*), characterized by a scale parameter *a* (see Supplementary Note 3, Table S1), the left-hand side of equation (2.6) depends on the dimensionless parameter 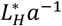 explicitly (see electronic supplementary material, Supplementary Note 4 for details). Therefore, for a landscape characterized by *A*_*H*_*A*^-1^, the equation (2.6) determines 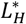 in units of the scale parameter *a* that characterises the pathogen dispersal kernel *b*(*x*). The threshold separation distance 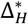 is then determined (on a regular square lattice, figure 1a) by 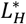 and *A*_*H*_*A*^-1^:

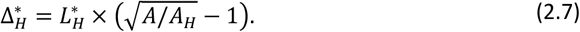

Parameters and functions used in the current paper are summarized in Table 1.

**Table 1.** Parameters used in the model, together with characteristic lengths of pathogen dispersal and host landscape and their threshold values.

| Symbol | Description |
| --- | --- |
| **Model parameters** |  |
| $A$ | Entire area of the landscape, a square plot of land. |
| $A_0$ | Area occupied by a single field of host crop (a square plot of cropped land). |
| $N$ | Number of host crop fields within the landscape. |
| $A_H$ | Total area occupied by fields (host crop),<br>$A_H = A_0 \times N$; $A_H \leq A$. |
| $n$ | Density of all host crop fields within a landscape, $n = N/A$. |
| $n_S$ | Density of susceptible fields within a landscape, assuming a single initial infected field,<br>$n_S = (N - 1)/A$. |
| $b(x)$ | Pathogen dispersal kernel. |
| $\sigma$ | Standard deviation of a Gaussian function. |
| $\beta$ | Infection rate per contact density. |
| $r$ | Rate of infection of susceptible fields at the start of an epidemic, equation (2.1). |
| **Spatial scales and degree of aggregation of a host landscape** |  |
| $a, b_0$ | Scale parameter ($a$) and a shape parameter ($b_0$) of a pathogen dispersal kernel, see Supplementary Material, Supplementary Note 3, Table S1. |
| $L_H, \Delta_H$ | Cluster size ($L_H$) and separation distance ($\Delta_H$) characterising a host crop landscape, figure 1a. |
| $N_{clusters}$ | Number of clusters of cropped fields within the landscape, $N_{clusters} = A_H/L_H^2 = A/(L_H + \Delta_H)^2$. It can be seen as a measure of the degree of aggregation of a host landscape. |
| **Threshold values** |  |
| $L_H^*, \Delta_H^*$<br>$N_{clusters}^*$ | Threshold values of cluster size ($L_H^*$) and separation distance ($\Delta_H^*$), figure 1c, expressed in units of the scale parameter $a$ and calculated from equations (2.6) and (2.7). $N_{clusters}^*$ is the corresponding threshold value of the number of clusters. |

### 2.4 Cassava landscape and CBSV

We illustrate the analytical approach with a real-life example: the invasion of CBSV in a cassava landscape. Godding et al. (2023)[31] developed and parameterised an epidemic spread model for CBSV with a pathogen dispersal kernel defined on a raster with a 1 km spatial resolution, imposed by the spatial resolution of available host distribution data [32]. The CBSV dispersal, determined in [31] by *β* and a function *K d*_*ij*_ of distance *d*_*ij*_ between centroids of raster cells, accounts for the combined effect of short-distance CBSV spread via insect vectors and long-distance spread via movement of virus-infected planting materials [31]. Specifically, *K d*_*ij*_ = 0 = *p* and *K d*_*ij*_ > 0 = *Cd*_*ij*_^-*a*^, where *p* is the proportion of inoculum remaining in the source cell, *C* is a normalization constant determined by ∑_*j*_ *K d*_*ij*_ = 1, and *α* is the kernel exponent. By fitting this model to surveillance data, Godding and co-authors [31] derived posterior distributions for the parameters *β, p* and *α*. Here, we use a radially symmetric, staircase-like version of the raster-based dispersal kernel from [31]. Using their results, we selected a power-law dispersal kernel *b*(*x*), characterised by two parameters: the exponent *α* = 3.75 and *p* = 0.12 taken from a region of relatively high probability density in posterior distributions estimated by Godding et al. (2023) [31].

In common with [31], we use CassavaMap [32] as a source of mapped landscape data for cassava. The landscape data for cassava are derived [31] from production statistics [32] and resolved to 1 km^2^ raster cells across sub-Saharan countries that cultivate cassava. The analytical estimates (2.6)-(2.7) depend on the ratio of host crop area *A*_*H*_ to the total area *A*. We express the landscape in terms of hectares of cassava fields per raster cell, noting that, according to [32], no more than half of the area of each raster cell (approximately 50 hectares) can be allocated to cassava production. Therefore, the maximum value of *A*_*H*_*A*^-1^ in the cassava landscape data cannot exceed 0.5, i.e. here *A*_*H*_*A*^-1^ ≤ 0.5.

Finally, using the approach described above, we calculated 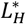 and 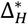 (figure 4a-b, black curves) for a cassava landscape at risk of CBSV invasion.

## 3. Results

Using the approximation for the rate *r* (equation 2.4, figure 2), we identified a range of values of cluster size *L*_*H*_ for which the infection rate, *r*, is minimized. Within this range, the largest cluster size, *L*_*H*_, is referred to as a threshold cluster size 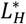. Equation (2.6) determines the ratio 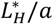 of the threshold cluster size 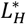 to the pathogen dispersal scale parameter *a* for a given pathogen dispersal kernel *b*(*x*) and the fraction *A*_*H*_*A*^-1^ of the landscape occupied by susceptible crops. Correspondingly, the threshold separation distance is calculated using equation (2.7).

### 3.1 Artificial landscapes and commonly-used dispersal kernels

Applying this approach to different dispersal kernels and landscapes, we first considered the Gaussian, negative exponential and power-law kernels – commonly used in epidemic spread studies [13,40,41] – alongside several additional kernels detailed in Supplementary Note 3, Table S1. The results for the exponential and power-law kernels are summarised in figure 3, while the results for the other kernels from Table S1 are presented in the electronic supplementary material, Supplementary Note 4.

**Fig. 3.**
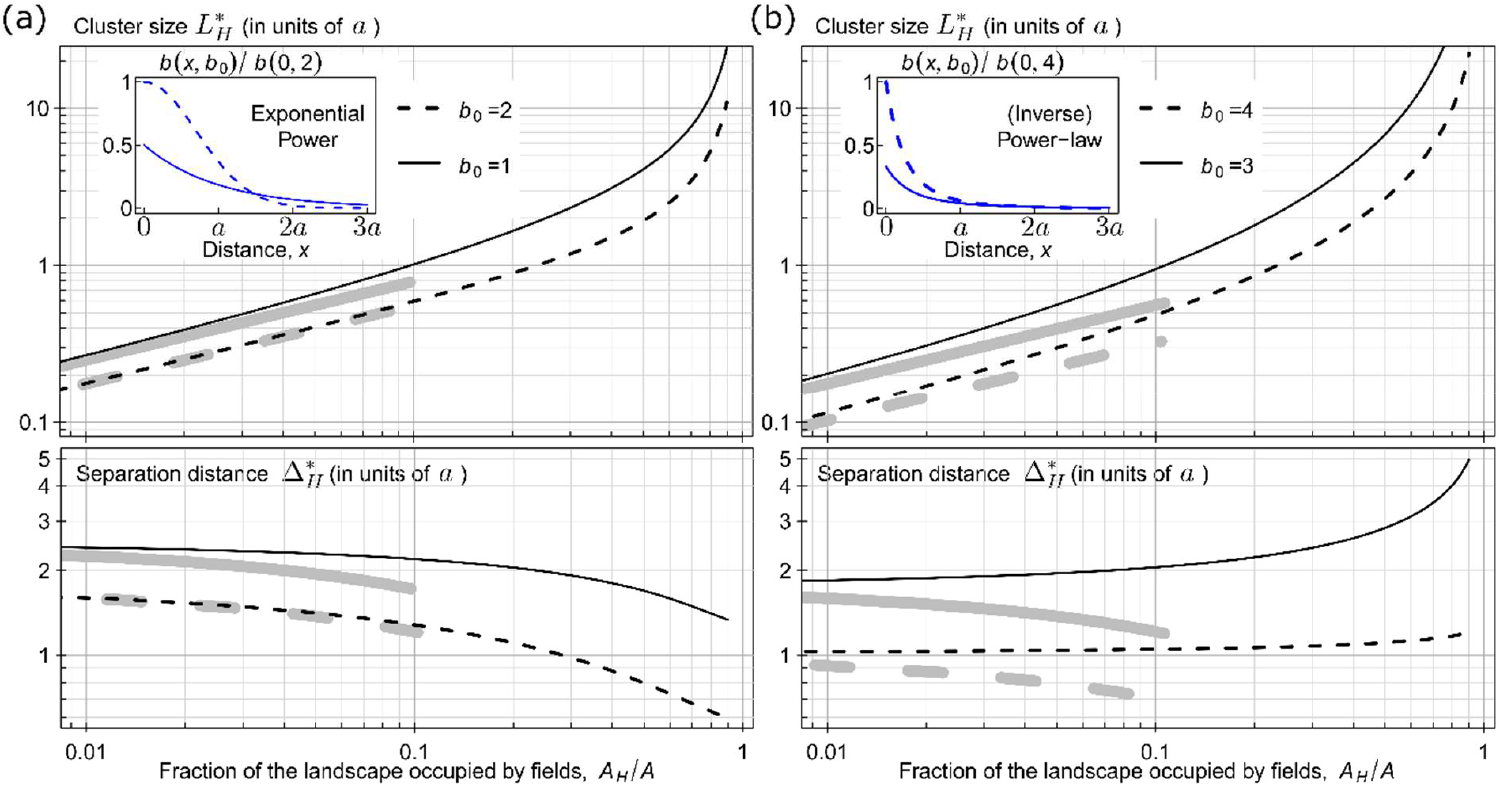
Threshold cluster size 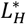 and threshold separation distance 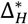 identified for two types of dispersal kernel. **(a)** Results are shown for exponential power kernels, that correspond to the Gaussian kernel when *b*_0_ = 2 and the negative exponential kernel when *b*_0_ = 1, as shown in the inset of the panel. **(b)** Results are shown for inverse power-law kernels with exponents *b*_0_ = 3 and *b*_0_ = 4, as illustrated in the inset of the panel. For the exact analytical expressions for the dispersal kernels used here see Table S1 in Supplementary Note 3. In both panels, black curves represent numerical estimates from equations (2.6) and (2.7), while grey curves at *A*_*H*_*A*^-1^ ≤ 0.1 represent explicit analytical estimates from equations (3.1) and (2.7). The R and Mathematica codes used to calculate these results are available from Figshare [25] and can be easily applied to other dispersal kernels.

In addition to the numerical solution for 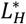 from equation (2.6), we derived an explicit analytical estimate for 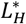 when the value of *A*_*H*_*A*^-1^ is small. In this case, the value of 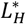is also small. Therefore, using the approximation 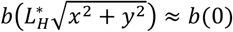 in equation (2.6) we derived:

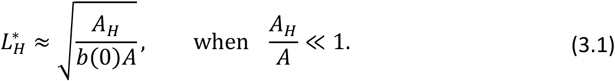

The explicit analytical estimate (equation 3.1) depends only on the dispersal kernel at zero distance, *b*(0), and ignores its distance-dependent component, *b*(*x* > 0). Consequently, the explicit analytical estimate of 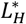 aligns better with the numerical solution for 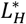 from equations (2.6–2.7) when short-range dispersal dominates, as with Gaussian kernels (figure 3a), but less so for long-tailed kernels like the power-law (figure 3b). This is reflected in the closer agreement between the thin black (for the numerical solution, equations 2.6–2.7) and the thick gray (explicit analytical estimate, equation 3.1) lines in figure 3a than in figure 3b. As the ratio *A*_*H*_/*A* increases, clusters move closer together, amplifying long-distance dispersal effects and reducing the accuracy of explicit analytical estimate (3.1), as seen in both panels.

We present an example of how results shown in figure 3 could inform stakeholders, including farmers and other agricultural planners, about the spatial structure of a host landscape. To keep the disease from spreading quickly at the start of an epidemic, it is important than no cluster of host crop fields is too large. Specifically, these clusters should not exceed a certain critical size (equal to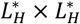 ). In addition, each cluster should be surrounded by a buffer zone – a strip of land of specific width (equal to 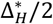) without susceptible host crops. Smaller clusters help limit disease spread within them, while buffer zones reduce the chance of transmission between clusters.

The same principle applies to clusters smaller than the critical size: any *L*_H_ × *L*_H_ clusters (where 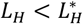) should be surrounded by a host-free buffer zone of width Δ_*H*_/2, where 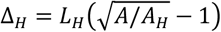 (cf equation (2.7)). The buffer zones around different clusters should not overlap. If there are two clusters within the landscape, first cluster 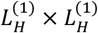 and second cluster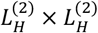, where both sizes are below the threshold size, i.e. 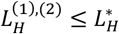, then the minimal suggested separation distance between these clusters equals the sum of the two corresponding buffer zones, i.e.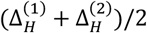.

Although we considered the spatial distribution of the cropped area into square clusters on a regular square lattice, other geometrical configurations could be considered. For example, if the cropped area is distributed among long narrow fields (e.g. see [8]), we present the derivation of an explicit analytical estimate for 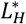 in Supplementary Note 5, analogous to equation (3.1).

### 3.2 Cassava landscape and an example of CBSV dispersal kernel

To demonstrate the application of the analytical approach to CBSV invasion in a cassava landscape, we used an area *A* = 24 × 24 km^2^ in an arbitrarily selected region on the border between Cameroon and the Central African Republic (figure 4c) derived from [32]. We calculated that cassava fields occupy 1.1% of this area (i.e. *A*_*H*_*A*^-1^ = 0.011). According to figures 4a and 4b, the corresponding threshold cluster size and separation distances are 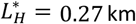 and 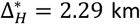. Therefore, if it were logistically possible, our analysis suggests that organizing the crop area from figure 4c into clusters of 0.27 km by 0.27 km 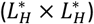, separated by 2.29 km 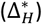, would result in the minimal infection rate, *r*_*min*_ = *βn*.

**Fig. 4.**
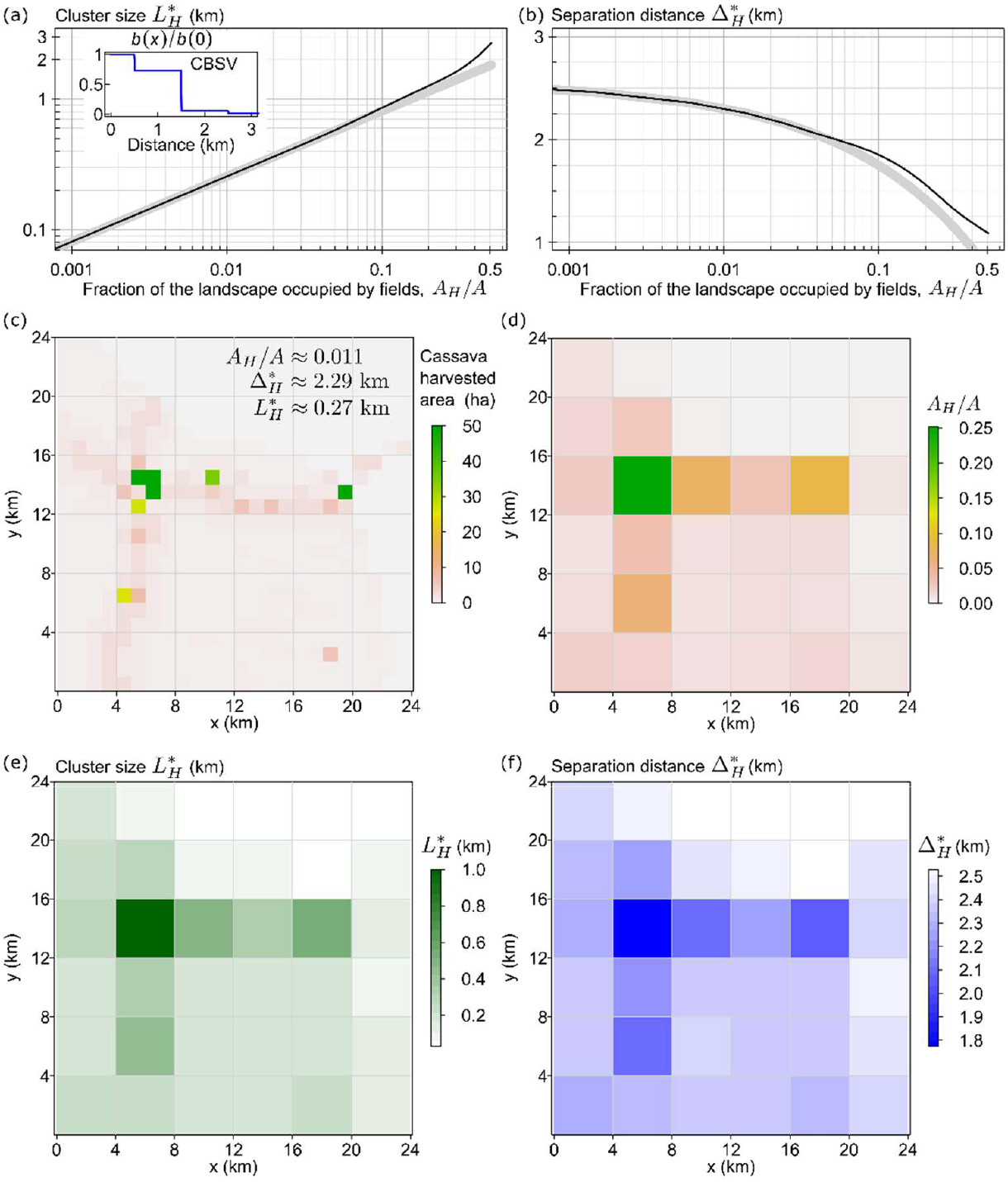
Threshold cluster size 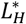 and threshold separation distance 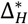 identified for an example of CBSV dispersal kernel and a cassava landscape. **(a)-(b)** Threshold cluster size 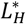 and separation distance 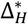 that minimise the initial infection rate *r* as the fraction *A*_*H*_/*A* of the landscape occupied by the host crop increases. Black curves represent numerical estimates obtained from equations (2.6) and (2.7), thick grey curves represent explicit analytical estimates from equations (3.1) and (2.7). **(c)** The cassava landscape, given at 1 km spatial resolution. **(d)** The same cassava landscape at 4 km spatial resolution, shown in terms of the fraction *A*_*H*_/*A* of area occupied by the host crop. **(e)-(f)** Maps of 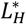 and 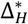 calculated using information from panels (a), (b) and (d), showing how the crop area in each cell can be maximally clustered for the infection rate to remain minimal (*r*_*min*_ = *βn*, with the value of *n* determined separately for each cell).

However, major redistributions of cropping patterns are unlikely to be practically feasible. Therefore, we considered a more practical approach by focusing on smaller 4 km by 4 km local areas where spatial reconfigurations of cropping patterns might potentially be possible. However, we emphasize that this is a hypothetical case to support the concept, as, in practice, there are many other major factors to be considered. We aggregated the original landscape to a 4 km spatial resolution (figure 4d). Using the fraction of area occupied by the host crop in each 4 km by 4 km cell, we assigned each cell a corresponding value of the threshold cluster size 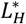 (figure 4e) and separation distance 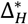 (figure 4f).

For illustration, consider the cell in figure 4d-f located between 4 km and 8 km on the x axis, and between 12 km and 16 km on the y axis. Our analysis indicates that the infection rate would remain minimal if cassava fields were aggregated into a cluster approximately 1 km by 1 km in size, surrounded by a cassava-free buffer zone with a width of 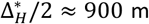. In another cell, located between 16 km and 20 km on the x axis and between 12 km and 16 m on the y axis, the infection rate would remain minimal if cassava fields were aggregated into a cluster of 500 m by 500 m, surrounded by a cassava-free buffer zone with a width of 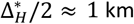.

Maps from figures 4e-f could in principle be used to inform decisions on where to plant new cassava fields without increasing the risk of early CBSV epidemic spread. If a cluster of size 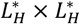 already exists within a given 4 km by 4 km cell, new fields should not be placed within the recommended cassava-free buffer zone (with a width of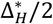) around the cluster to avoid increasing the infection rate. New fields can be planted near smaller clusters, but each smaller cluster of size 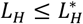 should be surrounded by a cassava-free buffer zone with a width of Δ_*H*_/2, where 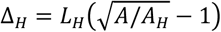.

## 4. Discussion

To identify the range of optimal clustering of host crops that minimizes the pathogen invasion rate, the analytical approach presented in this paper integrates the intrinsic spatial characteristics of pathogen dispersal with the distribution of host crops. Considering a specific geometrical configuration of host landscapes, where susceptible crops are distributed in square clusters on a regular square lattice, the resulting threshold characteristics, cluster size 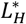 and separation distance 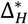, are derived in units of the scale parameter, *a*, of a given pathogen dispersal kernel and depend on the fraction *A*_*H*_/*A* of the landscape occupied by susceptible crops. Consequently, this analytical approach is effective for a wide range of dispersal kernels and host landscapes.

We used two main approximations in deriving the analytical results. The first approximation was applied to landscapes where hosts were aggregated into a *small number* of clusters (i.e. where *r* is a decreasing function of the number of clusters; see e.g. figure 2). In those cases, we estimated *r* from a single cluster and the corresponding density 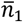 (that is the spatially averaged density of contacts with susceptible crops within a single cluster that can result in infection from an initial randomly infected host), 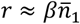, (approximation 1, equation (2.3)).

In other landscapes, where crops were aggregated into a *large number* of clusters (i.e. where *r* depends weakly on the number of clusters, see e.g. figure 2), we used a second approximation and estimated *r* assuming a constant density, *n* (that is the density of crops within the entire landscape and therefore independent of the number of clusters), *r ≈ βn*, (approximation 2).

Approximation 1 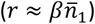 applies when clusters are sufficiently large and well-isolated, such that inter-cluster contacts are minimal. In this case, approximation 1 effectively considers each single cluster in isolation, neglecting inter-cluster contacts and therefore underestimating the actual value of *r*. In contrast, approximation 2 (*r ≈ βn*) applies when numerous sufficiently small clusters are located close to each other. In this case, contacts may occur across many clusters, except the most distant ones. However, because approximation 2 assumes contacts with all hosts across the entire landscape, it overestimates the actual value of *r*. The intersection of the two approximations determines the threshold cluster size 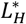, i.e. 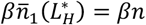 (equation 2.5), which is solved numerically to provide an exact value for 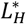. The underestimation (due to approximation 1) of the left-hand side of equation 2.5 and the overestimation on the right-hand side (due to approximation 2) lead to an underestimation at the intersection point for the thresholds number of clusters, 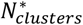 (figure 1c), and consequently, to an overestimation of 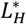 due to the relationship 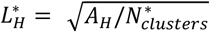 . The resulting overestimation of 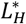 provides a practical upper bound to the size of a cluster of crop fields that minimizes the risk of rapid spread during the initial stages of epidemics.

By using methods based on *r* (or *R*_0_ given the link in SIR models as *R*_0_ = *r*/*μ* [24]), our analysis focuses on short-term epidemiological dynamics. The analysis is therefore useful for predicting outcomes during the initial invasion but does not necessarily provide an accurate assessment of the long-term dynamics of an epidemic. However, in some cases, medium-term dynamics can still be characterised by *r*. For example, in our earlier work [24] we demonstrated that “the analytical estimates of the infection rate correctly identified the relative ranking of the number of infected fields” [24] for different landscapes in sub-Saharan Africa, using a six-month spread period as a proxy for medium-term dynamics. In general, the nonlinear dynamics, typical of many epidemics, mean that initial fast or slow rates of epidemic progress are often not sustained throughout the epidemic. For example, Brown and Bolker [5] and Benincà et al. [42] have shown how initially high values of *r* or *R*_0_ can result in a slow spread of pathogens at later stages of an epidemic, where there are trade-offs between local and long-range dispersal. Therefore, to generate robust predictions for long-term epidemiological dynamics in realistic agricultural landscapes, it is necessary to use computer simulations of epidemic spread models. For instance, Godding et al. [43] modelled the 25-year spread of CBSV across sub-Saharan Africa, providing insights that go beyond what short-term analytical methods can offer.

The results of the current paper improve our understanding of the relationship between spatial scales of host distribution and pathogen dispersal, and their role in epidemic dynamics [7,44]. Specifically, the analytical results in figure 3 provide some general insights into the roles of the spatial scale parameter *a* and the shape parameter *b*_0_ (cf Table S1 in Supplementary Note 3) of a dispersal kernel in epidemics. These insights can be used to quickly compare how different dispersal kernels affect the invasion speed of pathogens in the same host landscape. This opens the door to future work on identifying optimal host distributions under invasion by multiple pathogens with different dispersal kernels. In addition, dispersal kernels are often poorly known, which makes the analytical approach presented here particularly valuable. It enables rapid exploration of a broad family of potential kernels – even before detailed empirical estimates are available.

For example, from equation (2.6) and its dependence on the dimensionless parameter 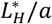, it follows that increasing the spatial scale parameter *a* (i.e., increasing dispersal length) reduces the infection rate at the start of an epidemic towards its minimal value *βn* (see Supplementary Note 6 for the details). Considering the role of the parameter *b*_0_, figure 3a shows that the values of 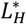 for the negative exponential kernel (*b*_0_ = 1) are larger than for Gaussian kernel (*b*_0_ = 2). This result indicates that, in the same host landscape, the infection rate *r* at the start of an epidemic will be higher for the Gaussian kernel than for the negative exponential kernel, assuming *β* is constant.

For ease of presentation, we have focused on pathogens, but the method can equally be applied to pest invasions where dispersal kernels are known or can be estimated. Ultimately, the analytical approach of this work has the potential to inform agronomic planning decisions [45,46] to optimise the size and spacing between clusters of fields, thereby minimizing the initial infection rate of an invading pathogen. The results of this paper provide valuable insights into selecting methods to decelerate the initial spread of an invading pathogen across a landscape of susceptible crops. Future work could further develop this analytical approach to encompass the role of characteristic spatial scales associated with control measures [7,23] on initial epidemic dynamics.

## Data availability

The datasets analysed during the current study were sourced from the previously published work by Suprunenko *et al*. [39] and are available in the Figshare repository, https://doi.org/10.6084/m9.figshare.25804702

## Acknowledgements

The authors are grateful to Dr Stephen Cornell for useful discussions at early stages of this work, and to Dr Alison Scott-Brown for helpful comments on the manuscript.

## Notes

**Funding** C.A.G. acknowledges financial support from the Gates Foundation (INV-010472).

### Competing Interest Statement

The authors have declared no competing interest.

### Summary of Updates

Title, Abstract, Introduction and Discussion revised, the key results remain unchanged. Supplementary Material updated to include Appendices from the earlier version of the manuscript.

