## Supplementary Notes 1-6. for "Optimizing crop clustering to minimize pathogen invasion in agriculture"

### Supplementary Material for "Optimizing crop clustering to minimize pathogen invasion in agriculture", Scientific Reports

Yevhen F. Suprunenko*, Christopher A. Gilligan

**Contents**

- Supplementary Note 1. Computer simulations of IBM.
- Supplementary Note 2. Derivation of the equation (2.3) from the main text.
- Supplementary Note 3. Dispersal kernels and corresponding threshold parameters.
- Supplementary Note 4. Dependence of $\overline{n}_1 A_0$ on dimensionless parameters.
- Supplementary Note 5. Approximate analytical results for long narrow fields. Derivation of the threshold parameters for long narrow fields.
- Supplementary Note 6. Effect of increasing spatial scale $a$ on infection rate $r$.

#### Supplementary Note 1. Computer simulations of IBM

The individual based model (IBM) from Suprunenko et al. (2025) (Ref. [24] in the main text), used in the current paper, is a stochastic, spatially-explicit IBM that describes the invasion and spread of a pathogen through a stationary population of identical square fields of crops within a square landscape. Each field is characterised by central coordinates and can be either susceptible or infected. Fields are aggregated into square clusters that are arranged spatially on a regular square lattice (figure 1a). Within a cluster, fields are arranged on a regular square lattice without overlap or gaps between individual fields.

At the start of an epidemic, a single randomly selected field within the entire landscape is infected. The dispersal of the pathogen is described by the product of the dispersal kernel, $b(x)$, and the epidemiological parameter $\beta$, that denotes the infection rate per contact density (figure 1b). Data (Ref. [39] in the main text) in figure 2 correspond to the following parameters used in computer simulations in Suprunenko et al. (2025) (Ref. [24] in the main text):

- Dispersal kernel $b(x) = \frac{1}{(2\pi\sigma^2)}\exp\left(-\frac{x^2}{2\sigma^2}\right)$, $\sigma = 10$ km and 1 km;
- Area of an individual field $A_0 = 0.04\,\mathrm{km}^2$;
- Total area of the entire landscapes $A = 365 \times 365\,\mathrm{km}^2$;
- Ratio of the area occupied by an agricultural crop, $A_H$, and the total area, $A_H/A = 0.6, 0.3, 0.15, 0.01$;
- Time is measured in units where the rate $\beta = 1$.

Suprunenko et al. (2025) (Ref. [24] in the main text) simulated epidemics using the 'Model simulator' software, developed by Cornell et al. (2019) (Ref. [37] in the main text) for continuous-time point process simulations. Using the expected total number of infected fields, $I(t)$, calculated as an average over 2000 simulations, the infection rate, $r$, was estimated as $r = \delta t^{-1} \times \ln[I(\delta t)/I(0)]$ using $\delta t = 0.1$.

### Supplementary Note 2. Derivation of the equation (2.3) from the main text

Using the definition of the quantity $\overline{n}_1$ presented in the main text,

$$\overline{n}_1 = \frac{\int\int b(|x_0 - x|)n_1(x_0)n_1(x)dx_0dx}{\int n_1(y)dy},$$

$\overline{n}_1$ can be expressed as:

$$\overline{n}_1 = \frac{1}{A_0}\frac{\int\int b(\sqrt{(x_0 - x)^2 + (y_0 - y)^2})f(x_0, y_0)f(x, y)dx_0dy_0dxdy}{\int f(x_0, y_0)dx_0dy_0},$$

where

$$f(x, y) = \begin{cases} 1, & x, y \in (0, L_H); \\ 0, & \text{otherwise.} \end{cases}$$

Using $\int f(x_0, y_0)dx_0dy_0 = L_H^2$ and the expression for $f(x_0, y_0)$ shown above, but keeping the function $f(x, y)$ and the integration over $x$ and $y$ over the entire domain, we obtain:

$$\overline{n}_1 = \frac{1}{A_0 L_H^2} \int_0^{L_H} dx_0 \int_0^{L_H} dy_0 \int_{-\infty}^{\infty} dx \int_{-\infty}^{\infty} dy b(\sqrt{(x_0 - x)^2 + (y_0 - y)^2}) f(x, y).$$

Next, change the variables of integration,

$$x = z_x + x_0;$$
$$y = z_y + y_0;$$

$$\overline{n}_1 = \frac{1}{A_0 L_H^2} \int_{-\infty}^{\infty} dz_x \int_{-\infty}^{\infty} dz_y b(\sqrt{z_x^2 + z_y^2}) \int_0^{L_H} dx_0 \int_0^{L_H} dy_0 f(z_x + x_0, z_y + y_0);$$

and measure all distances in units of $L_H$:

$$\overline{n}_1 = \frac{L_H^2}{A_0} \int_{-\infty}^{\infty} dz_x \int_{-\infty}^{\infty} dz_y b(L_H \sqrt{z_x^2 + z_y^2}) \int_0^1 dx_0 \int_0^1 dy_0 f(z_x + x_0, z_y + y_0).$$

The integral over $x_0$ and $y_0$ can be transformed into the following expression:

$$\int_0^1 dx_0 \int_0^1 dy_0 f(z_x + x_0, z_y + y_0) = \begin{cases} \int_{z_x}^1 dx_1 \int_{z_y}^1 dy_1, & 0 < z_x < 1, \quad 0 < z_y < 1 \\ \int_{z_x}^1 dx_1 \int_{-z_y}^1 dy_0, & 0 < z_x < 1, \quad -1 < z_y < 0 \\ \int_{-z_x}^1 dx_0 \int_{z_y}^1 dy_1, & -1 < z_x < 0, \quad 0 < z_y < 1 \\ \int_{-z_x}^1 dx_0 \int_{-z_y}^1 dy_0, & -1 < z_x < 0, \quad -1 < z_y < 0 \end{cases};$$

which becomes:

$$\int_0^1 dx_0 \int_0^1 dy_0 f(z_x + x_0, z_y + y_0) = \begin{cases} (1 - z_x)(1 - z_y), & 0 < z_x < 1, \quad 0 < z_y < 1 \\ (1 - z_x)(1 + z_y), & 0 < z_x < 1, \quad -1 < z_y < 0 \\ (1 + z_x)(1 - z_y), & -1 < z_x < 0, \quad 0 < z_y < 1 \\ (1 + z_x)(1 + z_y), & -1 < z_x < 0, \quad -1 < z_y < 0 \end{cases}.$$

Substituting the previous expression into the original expression for $\overline{n}_1$, we derive:

| Kernel name | Kernel $b(x)$ expression, $a > 0$ | Kernel name | Kernel $b(x)$ expression, $a > 0$ |
|---|---|---|---|
| (Negative) Exponential | $\frac{1}{2\pi a^2}\exp\left(-\frac{x}{a}\right)$ | Gaussian | $\frac{1}{\pi a^2}\exp\left(-\frac{x^2}{a^2}\right)$ |
| (Inverse) Power-law | $\frac{(b_0-2)(b_0-1)}{2\pi a^2}\times$ $\times\left(1+\frac{x}{a}\right)^{-b_0}$, $b_0 > 2$ | Exponential Power | $\frac{b_0}{2\pi a^2\Gamma\left(\frac{2}{b_0}\right)}\exp\left(-\frac{x^{b_0}}{a^{b_0}}\right)$, $b_0 > 0$ |
| 2Dt power-law | $\frac{(b_0-1)}{\pi a^2}\left(1+\frac{x^2}{a^2}\right)^{-b_0}$, $b_0 > 1$ | Tophat | $\begin{cases}\frac{1}{\pi a^2}, & x \le a\\ 0, & x > a.\end{cases}$ |
| Logistic | $\frac{b_0\left(1+x^{b_0}/a^{b_0}\right)^{-1}}{2\pi a^2\Gamma\left(\frac{2}{b_0}\right)\Gamma\left(1-\frac{2}{b_0}\right)}$, $b_0 > 2$ | | |

Table S 1: Dispersal kernels considered in this paper.

$$\overline{n}_1(L_H) = \frac{L_H^2}{A_0}\left[\int_0^1 dz_x \int_0^1 dz_y b(L_H\sqrt{z_x^2+z_y^2})(1-z_x)(1-z_y)+ \right.$$
$$+\int_0^1 dz_x \int_{-1}^0 dz_y b(L_H\sqrt{z_x^2+z_y^2})(1-z_x)(1+z_y)+$$
$$+\int_{-1}^0 dz_x \int_0^1 dz_y b(L_H\sqrt{z_x^2+z_y^2})(1+z_x)(1-z_y)+$$
$$\left.+\int_{-1}^0 dz_x \int_{-1}^0 dz_y b(L_H\sqrt{z_x^2+z_y^2})(1+z_x)(1+z_y)\right];$$
$$\overline{n}_1(L_H) = 4\frac{L_H^2}{A_0}\int_0^1 dz_x \int_0^1 dz_y b(L_H\sqrt{z_x^2+z_y^2})(1-z_x)(1-z_y).$$

This equation is the same as the equation (2.3) from the main text.

### Supplementary Note 3. Dispersal kernels and corresponding threshold parameters

We consider dispersal kernels that are widely used in phenomenological approaches in ecology and epidemiology, as summarized by Nathan et al. (2012) (Ref. [38] in the main text). Dispersal kernels $b(x)$ are normalized in 2D space, $\int_0^\infty \int_0^{2\pi} b(x)x\mathrm{d}x\mathrm{d}\phi = 1$, and include a scale parameter $a$ and a shape parameter $b_0$ (Table S1).

For each kernel in Table S1, we calculate threshold parameters $L_H^*$ and $\Delta_H^*$ using expressions (2.6) and (2.7) from the main text and the approximate estimate (3.1), i.e. $L_H^* \approx \sqrt{A_H A^{-1} b^{-1}(0)}$, also from the main text. Results are shown on Figure S1. The agreement between estimates (2.6) and (3.1) are better when kernels are shorter-ranged, for example: for Gaussian kernel the agreement is better than for negative exponential kernel, see Fig. 3(a) in the main text; for exponential power dispersal kernel with $b_0 = 3$ the estimates (2.6) and (3.1) are closer to each other than for the same kernel with $b_0 = 0.7$, see Fig. S1(a); for logistic kernel agreement between (2.6) and (3.1) is better for $b_0 = 4$ than for $b_0 = 2.3$, see Fig. S1(d). However, for inverse power-law dispersal kernel the agreement between (2.5) and (3.1) is of the same quality for both $b_0 = 4$ and $b_0 = 3$, see Fig. S1(b). For tophat dispersal kernel the estimates (2.6) and (3.1) are indistinguishable for small and medium values of the fraction $A_H/A$, see Fig. S1(e).

### Supplementary Note 4. Dependence of $\overline{n}_1 A_0$ on dimensionless parameters

Here we demonstrate that the expression $\overline{n}_1(L_H)A_0$ depends on the following dimensionless parameters: (i) $L_H/a$; (ii) a dimensionless parameter $b_0$ (if applicable). The explicit expressions for quantities $\overline{n}_1(L_H)A_0$ for each of the dispersal kernel from Table S1 are shown below. For brevity, the ratio $L_H/a$ is denoted by $w$:

$$w = \frac{L_H}{a}.$$

**Gaussian kernel**

$$b(x) = \frac{1}{\pi a^2}\exp\left(-\frac{x^2}{a^2}\right); \quad a > 0.$$

$$\overline{n}_1 A_0 = 4\frac{w^2}{\pi}\int_0^1 dz_x \int_0^1 dz_y \exp\left(-w^2(z_x^2 + z_y^2)\right)(1 - z_x)(1 - z_y).$$

**Negative exponential**

$$b(x) = \frac{1}{2\pi a^2}\exp\left(-\frac{x}{a}\right), \quad a > 0.$$

$$\overline{n}_1 A_0 = 2\frac{w^2}{\pi}\int_0^1 dz_x \int_0^1 dz_y \exp\left(-w\sqrt{z_x^2 + z_y^2}\right)(1 - z_x)(1 - z_y).$$

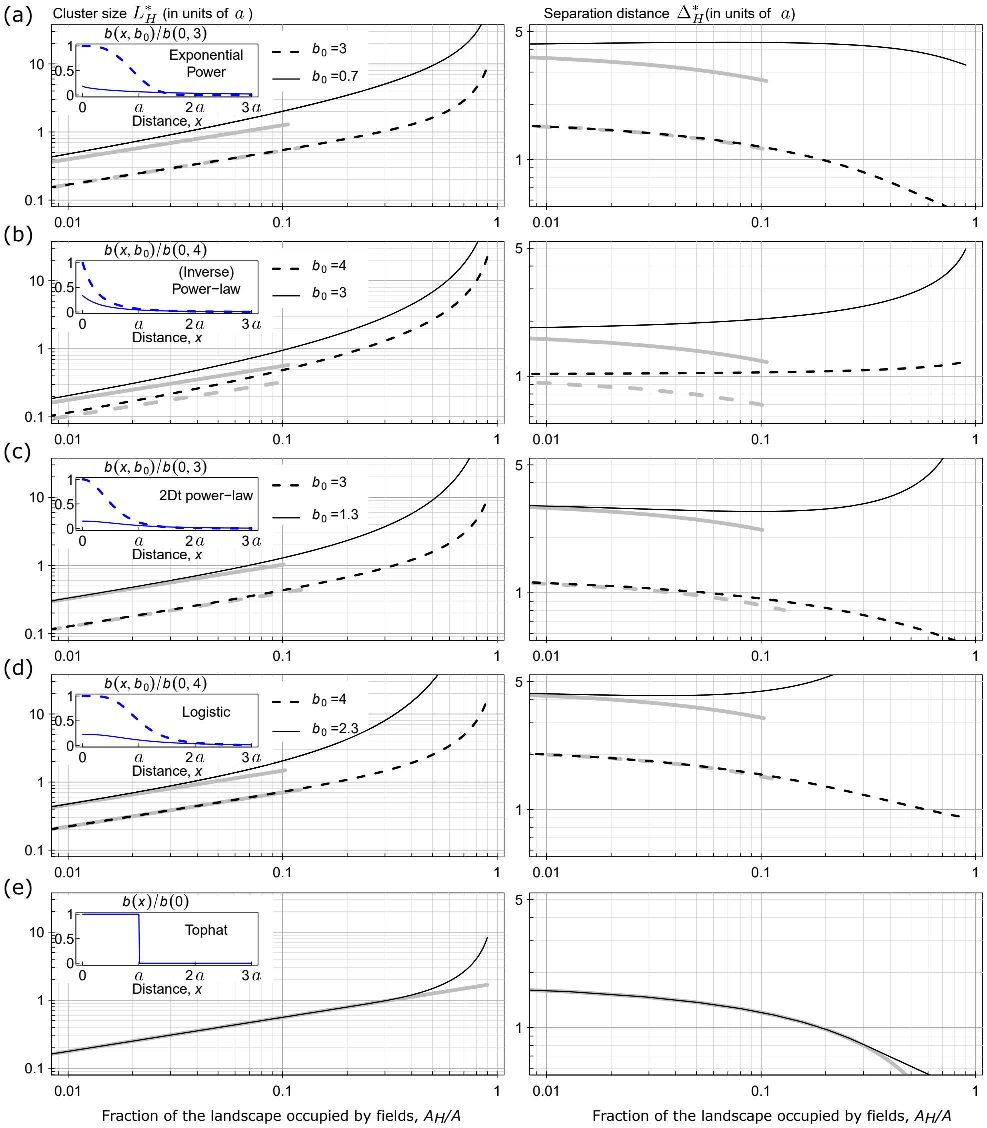


Figure S 1: The values of threshold parameters $L_H^*/a$ and $\Delta_H^*/a$ for dispersal kernels shown in Table S1. Each panel represents a row consistsing of two figures, $L_H^*/a$ as a function of $A_H/A$ on the left-hand side, and $\Delta_H^*/a$ as a function of $A_H/A$ on the right-hand side. All plots show an interval $A_H/A \in (0.08, 0.9)$. Black curves show estimates from equation (2.6) from the main text, thick gray curves show estimates based on equation (3.1) from the main text, i.e. $L_H^* \approx \sqrt{A_H A^{-1} b^{-1}(0)}$.

**Exponential Power (Gaussian for $b_0 = 2$; Exponential for $b_0 = 1$)**

$$b(x) = \frac{b_0}{2\pi a^2 \Gamma(2/b_0)} \exp\left(-\frac{x^{b_0}}{a^{b_0}}\right), \quad a, b_0 > 0.$$

$$\overline{n}_1 A_0 = 2\frac{w^2}{\pi}\frac{b_0}{\Gamma(2/b_0)} \int_0^1 dz_x \int_0^1 dz_y \exp\left(-w^{b_0}\left(z_x^2 + z_y^2\right)^{b_0/2}\right)(1 - z_x)(1 - z_y).$$

**2Dt power-law**

$$b(x) = \frac{(b_0 - 1)}{\pi a^2}\left(1 + \frac{x^2}{a^2}\right)^{-b_0}, \quad a > 0; b_0 > 1.$$

$$\overline{n}_1 A_0 = 4\frac{w^2}{\pi}(b_0 - 1) \int_0^1 dz_x \int_0^1 dz_y \left(1 + w^2\left(z_x^2 + z_y^2\right)\right)^{-b_0}(1 - z_x)(1 - z_y).$$

**(Inverse) Power-law**

$$b(x) = \frac{(b_0 - 2)(b_0 - 1)}{2\pi a^2}\left(1 + \frac{x}{a}\right)^{-b_0}, \quad a > 0; b_0 > 2.$$

$$\overline{n}_1 A_0 = 2\frac{w^2}{\pi}(b_0 - 2)(b_0 - 1) \int_0^1 dz_x \int_0^1 dz_y \left(1 + w\sqrt{z_x^2 + z_y^2}\right)^{-b_0}(1 - z_x)(1 - z_y).$$

**Logistic**

$$b(x) = \frac{b_0}{2\pi a^2 \Gamma\left(\frac{2}{b_0}\right)\Gamma\left(1 - \frac{2}{b_0}\right)}\left(1 + \frac{x^{b_0}}{a^{b_0}}\right)^{-1}, \quad a > 0; b_0 > 2.$$

$$\overline{n}_1 A_0 = 2\frac{w^2}{\pi}\frac{b_0}{\Gamma\left(\frac{2}{b_0}\right)\Gamma\left(1 - \frac{2}{b_0}\right)} \int_0^1 dz_x \int_0^1 dz_y \left(1 + w^{b_0}\left(z_x^2 + z_y^2\right)^{b_0/2}\right)^{-1}(1 - z_x)(1 - z_y).$$

**Tophat**

$$b(x) = \begin{cases} \frac{1}{\pi a^2}, & x \leq a; \\ 0, & x > a. \end{cases}$$

$$\overline{n}_1 A_0 = 4w^2 \int_0^1 dz_x \int_0^1 dz_y \frac{1}{\pi}\theta\left(1 - w\sqrt{z_x^2 + z_y^2}\right)(1 - z_x)(1 - z_y).$$

#### Supplementary Note 5. Approximate analytical results for long narrow fields. Derivation of the threshold parameters for long narrow fields

Here, we consider long narrow fields within a landscape of area $A$. The landscape contains the number $N_{fields}$ of long narrow fields, each with dimensions $l \times L_H$ where $l \gg L_H$. The fields are positioned along the $y$-axis and separated along the $x$-axis by a distance $\Delta_H$. The area occupied by fields is $A_H = N_{fields} \times l \times L_H$, and the total area of the landscape is $A = N_{fields} \times l \times (L_H + \Delta_H)$. Therefore, $\Delta_H = L_H \times (AA_H^{-1} - 1)$.

The equation that determines the threshold size $L_H^*$ of a field when $A_H A^{-1} \ll 1$, i.e. $L_H^* \sim A_H A^{-1} b^{-1/2}(0)$, is derived below.

Using the definition of the quantity $\overline{n}_1$ presented in the main text,

$$\overline{n}_1 = \frac{\int\int b(|x_0 - x|) n_1(x_0) n_1(x) dx_0 dx}{\int n_1(y) dy},$$

$\overline{n}_1$ can be expressed as:

$$\overline{n}_1 = \frac{1}{A_0} \frac{\int\int b(\sqrt{(x_0 - x)^2 + (y_0 - y)^2}) f(x_0, y_0) f(x, y) dx_0 dy_0 dx dy}{\int f(x_0, y_0) dx_0 dy_0},$$

where

$$f(x, y) = \begin{cases} 1, & x \in (0, L_H); \\ 0, & \text{otherwise.} \end{cases}$$

Note, the function $f(x, y)$ does not depend on the $y$ coordinate. Assume, the integral over $y$-coordinate is considered over the interval $y \in (-L_y/2, L_y/2)$ where $L_y \to \infty$. Using $\int f(x_0, y_0) dx_0 dy_0 = L_H L_y$ and the expression for $f(x_0, y_0)$ shown above, but keeping the function $f(x, y)$ and the integration over $x$ and $y$ over the entire domain, we obtain:

$$\overline{n}_1 = \frac{1}{A_0 L_H L_y} \int_0^{L_H} dx_0 \int_{-L_y/2}^{L_y/2} dy_0 \int_{-\infty}^{\infty} dx \int_{-\infty}^{\infty} dy b(\sqrt{(x_0 - x)^2 + (y_0 - y)^2}) f(x, y).$$

Next, change the variables of integration,

$$x = z_x + x_0;$$
$$y = z_y + y_0;$$

$$\overline{n}_1 = \frac{1}{A_0 L_H L_y} \int\limits_{-\infty}^{\infty} dz_x \int\limits_{-\infty}^{\infty} dz_y b(\sqrt{z_x^2 + z_y^2}) \int\limits_{0}^{L_H} dx_0 \int\limits_{-L_y/2}^{L_y/2} dy_0 f(z_x + x_0, z_y + y_0);$$

noticing that the integral over $y_0$ gives a pre-factor $L_y$, and measure all distances in units of $L_H$:

$$\overline{n}_1 = \frac{1}{A_0 L_H} \int\limits_{-\infty}^{\infty} dz_x \int\limits_{-\infty}^{\infty} dz_y b(\sqrt{z_x^2 + z_y^2}) \int\limits_{0}^{L_H} dx_0 f(z_x + x_0, ...);$$

$$\overline{n}_1 = \frac{L_H^2}{A_0} \int\limits_{-\infty}^{\infty} dz_x \int\limits_{-\infty}^{\infty} dz_y b(L_H \sqrt{z_x^2 + z_y^2}) \int\limits_{0}^{1} dx_0 f(z_x + x_0, ...).$$

The integral over $x_0$ can be transformed into the following expression:

$$\int\limits_{0}^{1} dx_0 f(z_x + x_0, ...) = \begin{cases} \int_{z_x}^{1} dx_1, & 0 < z_x < 1, \\ \int_{-z_x}^{1} dx_0, & -1 < z_x < 0, \end{cases} ;$$

which becomes:

$$\int\limits_{0}^{1} dx_0 f(z_x + x_0, ...) = \begin{cases} (1 - z_x), & 0 < z_x < 1, \\ (1 + z_x), & -1 < z_x < 0, \end{cases} .$$

Substituting the previous expression into the original expression for $\overline{n}_1$, we derive:

$$\overline{n}_1(L_H) = \frac{L_H^2}{A_0} \left[ \int\limits_{0}^{1} dz_x \int\limits_{-\infty}^{\infty} dz_y b(L_H \sqrt{z_x^2 + z_y^2})(1 - z_x) + \right.$$

$$\left. + \int\limits_{-1}^{0} dz_x \int\limits_{-\infty}^{\infty} dz_y b(L_H \sqrt{z_x^2 + z_y^2})(1 + z_x) \right] ;$$

$$\overline{n}_1(L_H) = 4 \frac{L_H^2}{A_0} \int\limits_{0}^{1} dz_x \int\limits_{0}^{\infty} dz_y b(L_H \sqrt{z_x^2 + z_y^2})(1 - z_x).$$

Asuming $L_H << 1$, we obtain:

$$\overline{n}_1(L_H) = 4\frac{L_H^2}{A_0}\int_0^1 dz_x(1-z_x)\int_0^\infty dz_y b(L_H\sqrt{z_y^2})$$
$$= 2\frac{L_H}{A_0}\int_0^\infty dz b(z)$$

The integral can be calculated explicitly for dispersal kernels from Table S1. In all cases, the following relationship holds (as shown in the file code.nb in the code used in this work available online from Ref. [25] in the main text):

$$\overline{n}_1(L_H) \sim \frac{L_H}{A_0}\sqrt{b(0)}$$

Therefore, the threshold value $L_H^*$ is determined by the condition $\overline{n}_1(L_H^*)A_0 = A_H A^{-1}$, that becomes:

$$L_H^* \sim \frac{1}{\sqrt{b(0)}}\frac{A_H}{A}.$$

### Supplementary Note 6. Effect of increasing spatial scale $a$ on infection rate $r$

Consider one of the kernels shown in figure 3 and a landscape area with a fixed value of $A_H A^{-1}$. Denote the corresponding value of $L_H^* a^{-1}$ as $L_0$. For two different values of the scale parameter, $a_1$ and $a_2$, we have $L_H^{*(1)} = L_0 \times a_1$, and $L_H^{*(2)} = L_0 \times a_2$. Therefore, for $a_1 > a_2$, we have $L_H^{*(1)} > L_H^{*(2)}$.

These threshold sizes, $L_H^{*(1)}$ and $L_H^{*(2)}$, correspond to the same value of infection rate, $r_{min} = \beta n$, where $n = A_H A^{-1} A_0^{-1}$. Denote the infection rates corresponding to scale parameters $a_1$ and $a_2$ as $r^{(1)}$ and $r^{(2)}$, respectively, and consider them in the following two configurations of the given landscape:

1. Distributing the given cropped area $A_H$ into clusters of a smaller size, $L_H^{*(2)}$, results in both infection rates being equal to the minimal rate, $r^{(1)} = r^{(2)} = r_{min}$.

2. Redistributing the cropped area $A_H$ into clusters with a larger size, $L_H^{*(1)}$, does not change the infection rate $r^{(1)}$, i.e. $r^{(1)} = r_{min}$, but increases $r^{(2)}$ because $L_H^{*(1)} > L_H^{*(2)}$.

Hence, for $a_2 < a_1$ we would get $r^{(2)} \geq r^{(1)} \geq r_{min}$. In other words, increasing the scale $a$ from $a_2$ to $a_1$ decreases the infection rate of pathogen dispersal towards its minimal value $r_{min}$.
